# Proteogenomic single cell analysis of skeletal muscle myocytes

**DOI:** 10.1101/2020.01.23.916791

**Authors:** Katherine M. Fomchenko, Rohan X. Verma, Suraj Kannan, Brian L. Lin, Xiaoping Yang, Tim O. Nieuwenhuis, Arun H. Patil, Karen Fox-Talbot, Matthew N. McCall, Chulan Kwon, David A. Kass, Avi Z. Rosenberg, Marc K. Halushka

## Abstract

Skeletal muscle myocytes have evolved into slow and fast-twitch types. These types are functionally distinct as a result of differential gene and protein expression. However, an understanding of the complexity of gene and protein variation between myofibers is unknown. We performed deep, whole cell, single cell RNA-seq on intact and fragments of skeletal myocytes from the mouse flexor digitorum brevis muscle. We compared the genomic expression data of 171 of these cells with two human proteomic datasets. The first was a spatial proteomics survey of mosaic patterns of protein expression utilizing the Human Protein Atlas (HPA) and the HPASubC tool. The second was a mass-spectrometry (MS) derived proteomic dataset of single human muscle fibers. Immunohistochemistry and RNA-ISH were used to understand variable expression. scRNA-seq identified three distinct clusters of myocytes (a slow/fast 2A cluster and two fast 2X clusters). Utilizing 1,605 mosaic patterned proteins from visual proteomics, and 596 differentially expressed proteins by MS methods, we explore this fast 2X division. Only 36 genes/proteins had variable expression across all three studies, of which nine are newly described as variable between fast/slow twitch myofibers. An additional 414 genes/proteins were identified as variable by two methods. Immunohistochemistry and RNA-ISH generally validated variable expression across methods presumably due to species-related differences. In this first integrated proteogenomic analysis of mature skeletal muscle myocytes we confirm the main fiber types and greatly expand the known repertoire of twitch-type specific genes/proteins. We also demonstrate the importance of integrating genomic and proteomic datasets.

## Introduction

Skeletal muscle is a voluntary, striated muscle found throughout the body with contraction regulated by nerve impulses through the neuromuscular junction (NMJ). Skeletal muscles consist of different fiber types delineated by the isoform of the myosin heavy chain they express, metabolic function, and other properties (1). In humans, slow fibers (type 1) and some fast fibers (type 2A) exhibit oxidative metabolic properties, while fast type 2X fibers exhibit glycolytic metabolic properties (2). Mice have an additional type 2B fast fiber. These fiber types are variable across different muscles of the body reflecting different functional needs (2, 3).

Multiple proteins and protein classes vary across fiber types (1, 4). These include isoforms of the myosin heavy and light chains, calcium ATPase pumps, troponin T, and tropomyosin proteins, as well as metabolic proteins, such as pyruvate kinase, GAP dehydrogenase, and succinate dehydrogenase. Beyond these classes, there have been few efforts to catalog the entirety of fast/slow twitch expression differences by proteomics or genomics.

Among proteins, the deepest effort, to date, has been the single fiber proteomics work of the Mann laboratory (5, 6). In separate studies of mouse and human single fiber skeletal muscles, 1,723 and 3,585 proteins were reported, respectively, many of which were variably expressed among slow and fast twitch fibers. The most comprehensive gene expression study was performed in mice using DNA microarrays across ten type 1 and ten type 2B fibers (7). Single cell RNA-sequencing (scRNA-seq) also has been performed in skeletal muscle and muscle cultures. However, the large size of skeletal myocytes has precluded them from these datasets, which are instead predominately satellite cells, and other supporting cell types (8–15). A recent publication used SMART-Seq to evaluate three fast twitch mouse fibers (16). The totality of these studies strongly suggests there are numerous expression differences between skeletal muscle fiber types and a need for new approaches to capture this diversity.

The Kwon laboratory recently developed a large cell sorting method to isolate mature cardiac myocytes (17). We ascertained if this method could be used to isolate the even larger skeletal muscle myocytes for scRNA-seq. Our goal was to combine this genetic data with single cell spatial proteomic data from the Human Protein Atlas (HPA) and an established mass spectrometry human skeletal muscle proteomic dataset for a unique proteogenomic characterization of skeletal muscle expression mosaicism.

## Results

### scRNA-seq and identification of fast/slow twitch fiber types

We performed single cell RNA-seq using the established mcSCRB-seq protocol (18, 19). We recovered data for 763 cells and sequenced to a median depth of 108,110 reads per cell. As we were unsure of where the ideal skeletal myocytes might arise from our flow-sorting method, they were taken from two different gates set on extinction (EXT) always “high” and time of flight (TOF) being both high or low (Supplementary Fig. 1). Additional cells were collected from a pseudo-biopsy approach with fragmented skeletal myocytes (see methods). Preliminary analyses, however, indicated a distinct cluster of cells with a high percentage of mitochondrial reads or otherwise low abundance reads. Notably, almost all of our pseudo-biopsy myocyte fragments and many TOF-low cells fell into this category. These quality control metrics likely indicated poor quality or sheared cells with loss of RNA. Thus, we excluded these cells and narrowed our analysis to the best 171 cells remaining with a median read count of 239,252 per cell.

An average of 12,098 transcripts were identified in these cells and all had the expression patterns of mature skeletal myocytes, highly expressing a myosin heavy chain isoform. Because of the narrow focus of this work to delineate cell subtypes and expression variability of just skeletal muscle myocytes, this isolation strategy linked to deep sequencing, proved to be advantageous.

We performed PCA of the data, corrected the data for the top 20 PCAs and utilized the top 3,000 variable genes (by +/− standard deviation) to cluster these cell types (Fig. 1a). Three groups were observed in a UMAP dimensionality reduction plot. The first cluster, containing 69 cells (40% of all cells) had elevated expression of *Myh1* and *Myh8* clearly identifying this group as containing fast 2X type cells and denoted as fast 2X_a_ (Fig. 1d). A second cluster (N=53 cells) had slightly more variable *Myh1* and *Myh8* differential expression, but by overall *Myh* gene expression, Fig. 1g, also appeared to be a fast 2X cell type (denoted fast 2X_b_). Of note, *Myh4*, a myosin heavy chain associated with fiber type 2B, was elevated in a single cell in this group (Fig. 1g) (3).

**Supplementary Figure 1.**
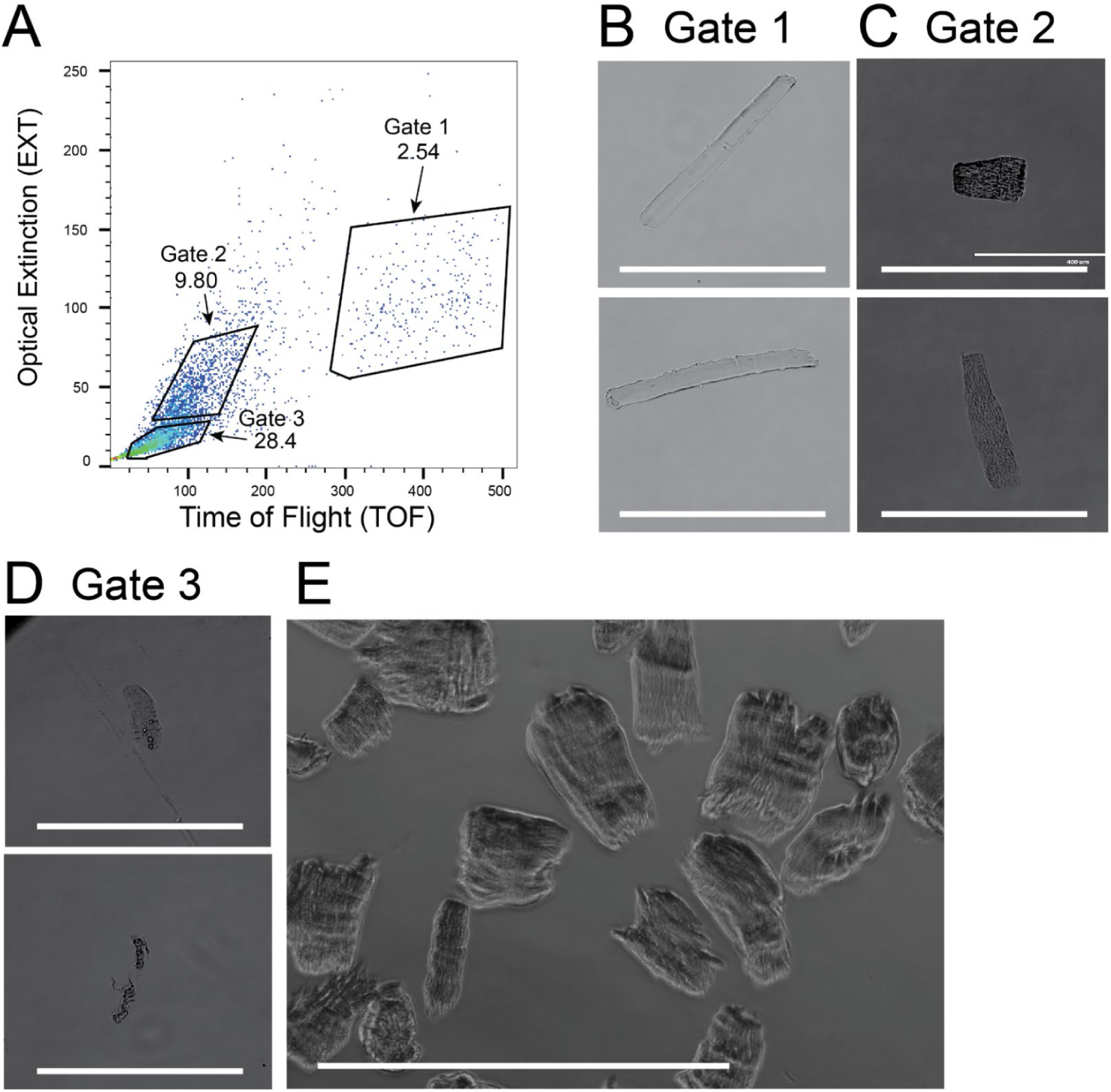
Mouse skeletal muscle myocyte preparation. A) Flow cytometry showing three gated areas representing EXT-high/TOF-high, EXT-high/TOF-low and EXT-low populations of flexor digitorum brevis myocytes. B) Representative images of Gate 1 EXT-high/TOF-high. C) Representative images of Gate 2 EXT-high/TOF-low. D) Representative images of Gate 3 EXT-low. E) Representative image of pseudo-biopsy isolated myocyte fragments. Gates 1 and 2 were used for library preparation. White size bar is 400 µm.

**Fig. 1.**
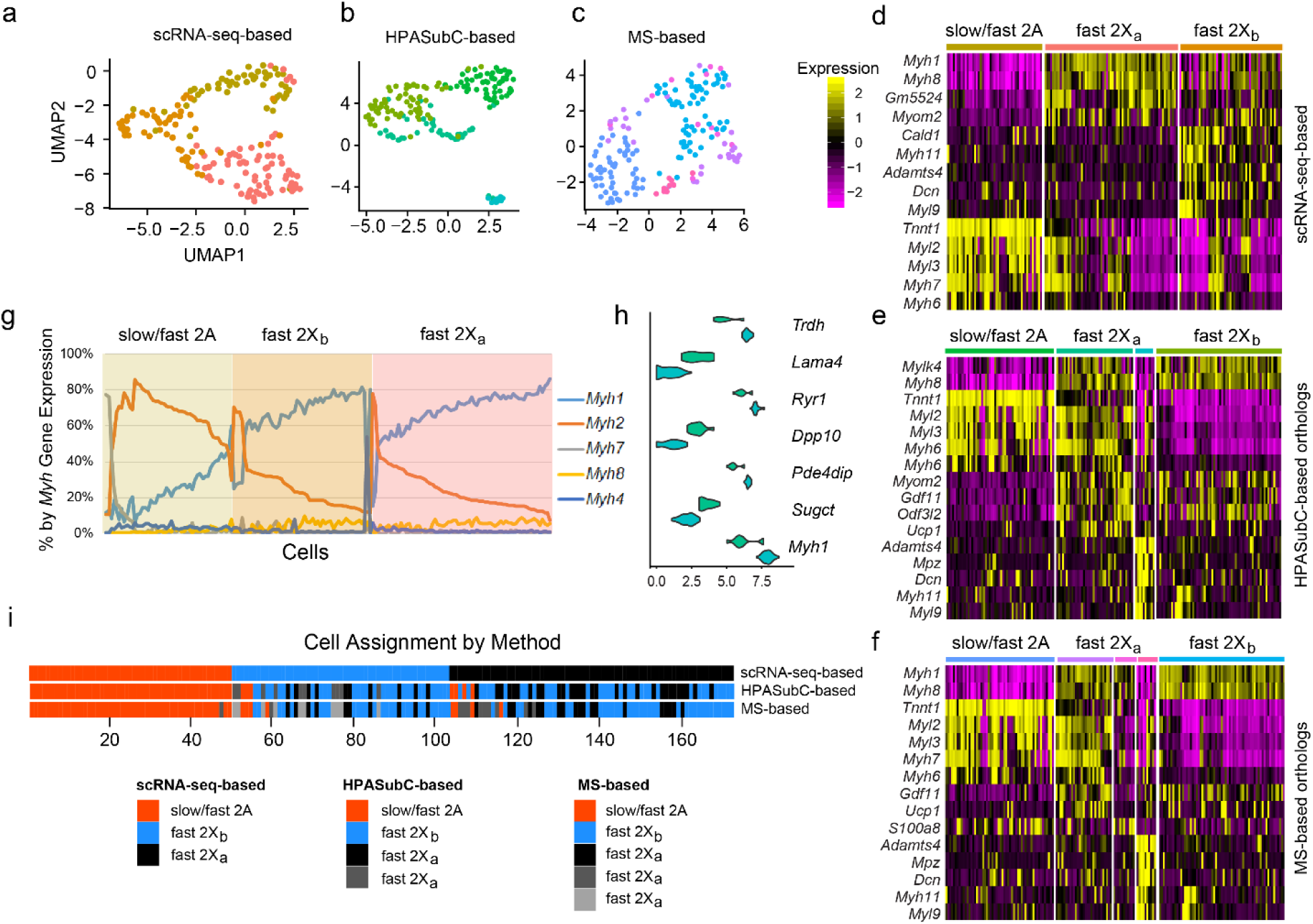
**a)** UMAP graph of 171 skeletal muscle cells based on variable gene expression determined by scRNA-seq. **b)** UMAP graph based on mouse orthologous expression of HPASubC variable proteins. **c)** UMAP graph based on mouse orthologous expression of MS variable proteins **d-f)** Heat maps of major genes expression differences between the different fiber types based on the different datasets. **g)** Major myosin heavy chain distributions across the 171 cells as a percentage of each heavy chain. The colored areas are the assignments of each cell based on the scRNA-seq-based data **h)** Violin plots of 7 genes that varied between the two fast 2X_a_ groups in the HPASubC-based data set. **i)** Assignment of each skeletal myocyte to a fiber type across the three methods. Strong agreement existed for the slow/fast 2A cells by any method of analysis

A third cluster (3) containing 49 cells (29% of the total) was defined by high expression of *Tnnt1* and *Myh2.* A deeper analysis of this group showed that 12 cells had high to modestly elevated *Myh7* expression (a slow-twitch marker), indicating this cluster was a combination of slow-twitch cells and fast 2A fibers (Fig. 1g). The flexor digitorum brevis is a fast twitch muscle, thus the overall distribution of significantly more fast (159) to slow fibers (12) is consistent with expected.

Interestingly, the expression patterns of the main fast/slow differentiating *Myh* genes was not as dichotomous as noted in protein based fiber type data (5). Here there were many more cells with intermediate levels and coexpression of *Myh1* and *Myh2* suggesting higher gene plasticity and more cell hybrids (Fig. 1g) (3).

### HPA-based mosaic protein discovery

To complement variable gene expression data, we generated mosaic protein data by performing an analysis of the IHC-based HPA dataset of skeletal muscle images using the HPASubC suite of tools (20). The HPASubC tool, obtains a selected organs’ images from the Human Protein Atlas (HPA) and allows rapid and agnostic interrogation of images for staining patterns of interest. This approach established a protein-based list of mosaically-expressed proteins. Out of 50,351 images reviewed for 10,301 unique proteins, 2,164 proteins had possible mosaic expression in skeletal muscle. Based on the aggregate image scores assigned to each protein, they were subsetted into categories of “real” mosaicism (374 proteins), “likely” mosaicism (1,231 proteins), and “unknown” probability of mosaicism (559 proteins) (Supplementary Data 1, Supplementary Fig. 2). For analysis purposes, we focused on the 1,605 proteins that were in the “real” or “likely” categories to reduce the incidence of false positive staining.

This method identified the well-known fiber type specific proteins such as MYH1, MYH2, MY4, MHY6, MHY7, and MYH8 that were categorized as both “real” or “likely” based on staining patterns (Supplementary Data 1). It also identified numerous uncharacterized or poorly characterized proteins, such as the zinc finger proteins ZNF213, ZNF282, ZNF343, ZNF350 and ZNF367 all of which had “real” patterns of mosaicism. A limitation of this spatial, IHC-based approach is that each protein image is independent of other proteins. Thus, one cannot identify co-expression patterns to assign proteins to certain fiber types.

**Supplementary Figure 2.**
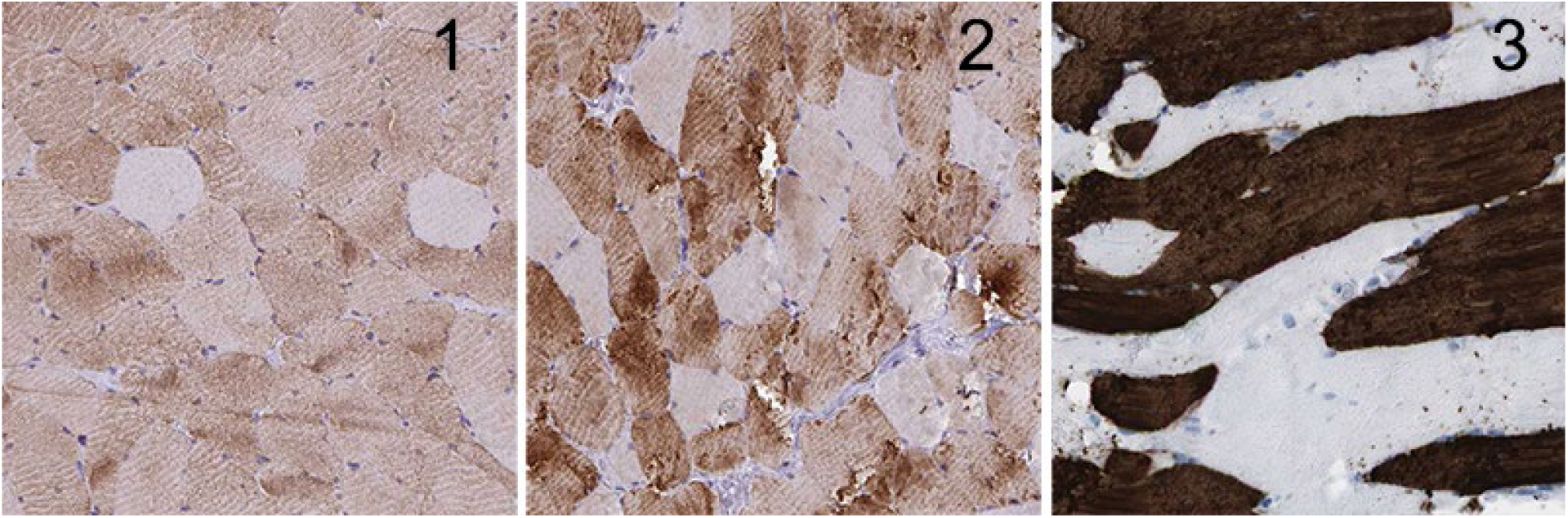
Scoring schema for HPASubC-based skeletal muscle mosaicism. A score of 1 indicated an “unknown” mosaicism based on subtle differences in stain intensity, or inconsistent patterns. A score of 2, “likely,” was a clear distinction of staining by myofiber but the staining was not robust. A score of 3 “real” identified clear and robust staining differences by muscle cell. The score was primarily about the pattern and secondarily about the intensity of the staining difference.

We therefore investigated how these 1,605 proteins might inform on fiber type of skeletal muscle cells by using this list to subset the orthologous mouse gene data from the scRNA-seq experiment. Using just these orthologous mouse genes, we regenerated the UMAP plot that identified four clusters (Fig. 1b). It essentially recapitulated the fast and slow fibers types noted from the exclusive scRNA-seq data, despite being based on a different set of genes. Uniquely, it subsetted the fast 2X_a_ cluster into two groups, one denoted by high expression of *Myom2* and *Gdf11* and the other denoted by high *Ucp1* and *Adamts4* (Fig. 1e). A t-test of gene expression comparing genes from just these two subsets of the fast 2X_a_ cluster identified multiple genes variably expressed between them (Fig. 1h). Although the cell clustering was generally similar between mouse scRNA-seq gene data and HPASubC data with regard to slow/fast 2A vs fast 2X, it was unclear which method was more representative. Therefore, we obtained a public MS dataset as a third method to classify slow and fast twitch fibers.

### Fast/slow twitch variation by MS-based proteomics

The human skeletal muscle fiber MS data in Murgia et al. is based on 152 fibers from eight donors (5). This dataset had 596 proteins with >2.3 fold variation between type 1 and type 2A fibers. We analyzed the full LFQ dataset of protein expression and constructed a UMAP plot that showed four clusters (Supplementary Fig. 3). One cluster was composed primarily of slow type 1 fibers and was adjacent to a second cluster with a small mixture of slow and other cell types. Two other clusters were primarily a collection of fast 2X and fast 2A cell types. Similar to the HPASubC approach above, we subsetted the orthologous mouse genes to these 596 proteins to explore cell fiber type assignment.

**Supplementary Figure 3.**
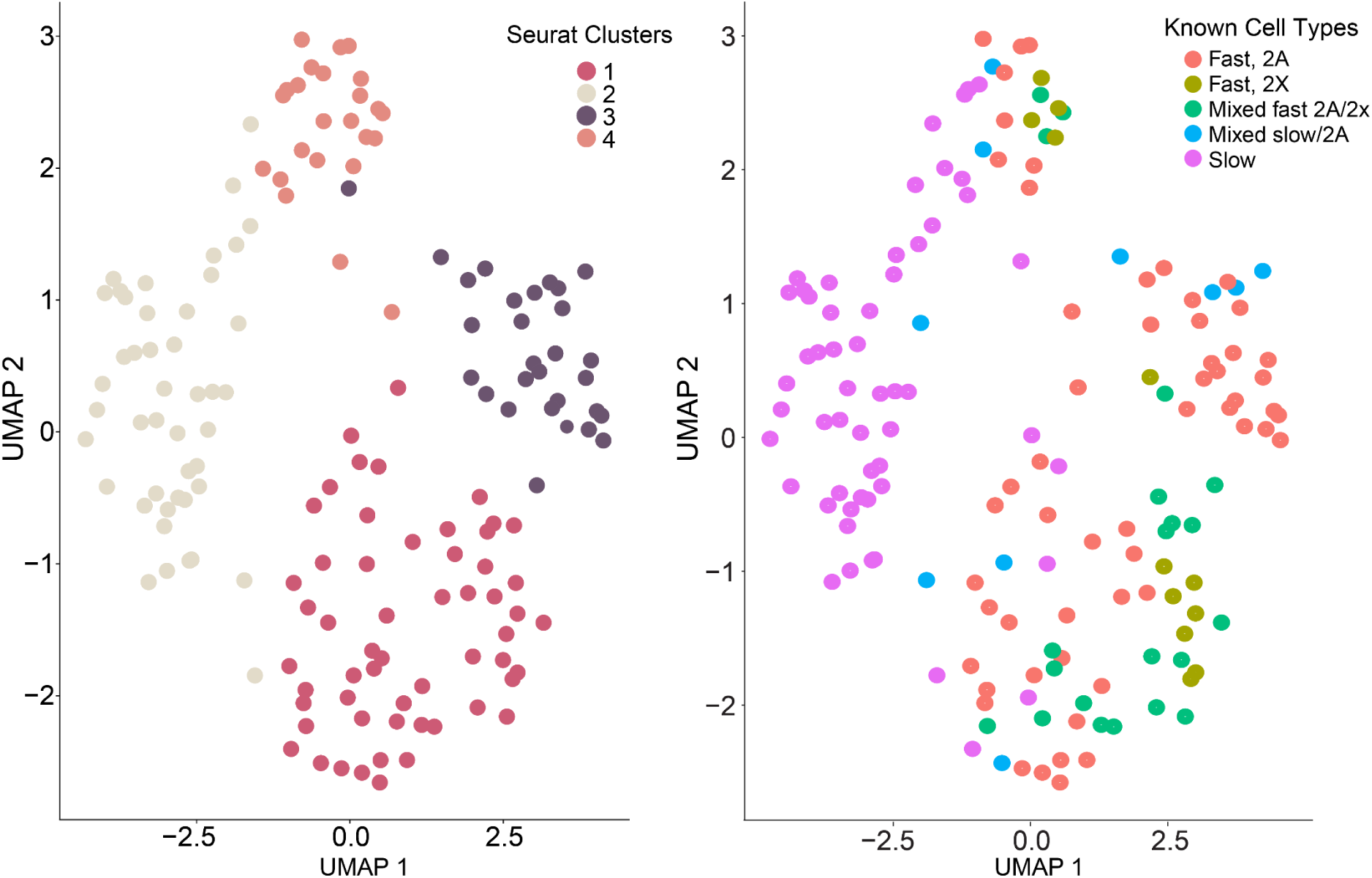
UMAP of MS-based protein data by cell type. **A)** Seurat identified four cell clusters. **B)** UMAP was coloured based on cell assignments of Murgia et al. The slow type cells are generally Seurat cluster 2. Fast 2A cells are generally in Seurat cluster 3, although they are also detected in clusters 1 and 3. Fast 2X clusters are predominately in Seurat cluster 1.

As seen in the UMAP plot, five groups were identified (Fig. 1c). Similar to the other two datasets (scRNA-seq and HPASubC), a slow/fast 2A fiber type was denoted by elevated expression of several genes including *Tnnt1* and *Myl2* (Fig. 1f). One fast 2X fiber group (2X_b_) was identified by high expression of *Myh1* and *Myh8.* The second fast 2X fiber group was then subdivided into three groups based on alternative elevated expression of genes that include *Gdf11* and *Ucp1* (group 3), *S100A8* (group 4) and *Adamts4* and *Mpz* (group 5). Unlike the protein expression level based UMAP, slow fibers and fast 2A fibers were not distinct. (Supplementary Fig. 3). This difference may be a result of the higher percentage of slow fibers in the MS dataset.

### Cross comparisons of the three approaches yield similar cell types

We identified the cluster assignment of each skeletal muscle cell based on the scRNA-seq, HPASubC, and MS approaches. We then plotted this information to demonstrate the extent to which there was fluidity in assignment by fiber type (Fig. 1i). All but one cell (48/49) assigned to the slow/fast 2A cluster based on scRNA-seq data remained in that cluster using other methods of clustering (HPASubC and MS). An additional 7-8 cells from the fast 2X groups became assigned to the slow/fast 2A cluster using the other methods of cell assignment. Cells moved interchangeably between the fast 2X_a_ and fast 2X_b_ clusters depending on the method used to cluster. We used this information to try and understand what distinguished fast 2X_a_ and fast 2X_b_ clusters.

### The 2X_a_ and fast 2X_b_ clusters differ by axonal genes

To understand if the two fast 2X clusters represent unique cell types, cell states, or some technical division, we performed a differential expression to determine what genes drove their differences. Of 5,260 genes compared, 557 genes were differentially expressed (t. test; adj. p. value <0.01). A Gene Ontology (GO) analysis on the 557 genes identified an enrichment of the cellular component “neuronal synapse,” suggesting variability at the NMJ. A further review of the top significant genes showed that >20 genes appear to have neuronal origins (*Cdh4, Cdkl5, Cntn4, Dscam, Gabbr2, Kirrel3, Lingo2, Lrp1, L1cam, Nrcam, Ntn1, Ntrk3, Ptprt, Ptpro, Robo2, Sdk1, Sema5a, Sema6d, Shank2, Sox5, Tnr,* and *Wwox*). Of these, NRTK3, LRP1, and ROBO2 were identified as mosaic in skeletal muscle cells by HPASubC. Additionally, in HPA images, seven orthologous proteins of these “neuronal” genes showed moderate staining, but each of these had a TPM <1 (from GTEx expression data). Only LRP1 was identified in the orthologous MS-dataset. This variability made us wonder how frequently the same genes/proteins were noted to be mosaic by each of the three methods.

### There is limited overlap of shared expression information

We compared the 3,000 most variable genes, the 1,605 HPASubC proteins, and the 596 MS proteins for shared patterns of mosaicism (Fig. 2). Only 23 genes/proteins were mosaic by all three approaches using the Seurat analysis method (Fig. 2a). An additional 300 genes/proteins were shared across two methods, with the most overlap identified between the two datasets with the most genes/proteins. Thus, we reasoned the abundance of genes/proteins by method was a major driver of overlap leading us to focus on the 3,052 transcripts shared by all three approaches, regardless of their mosaic/variable status. This resulted in 157 genes/proteins shared across any two methods with the most overlap between the two protein datasets (77 proteins) (Fig. 2c).

**Fig. 2.**
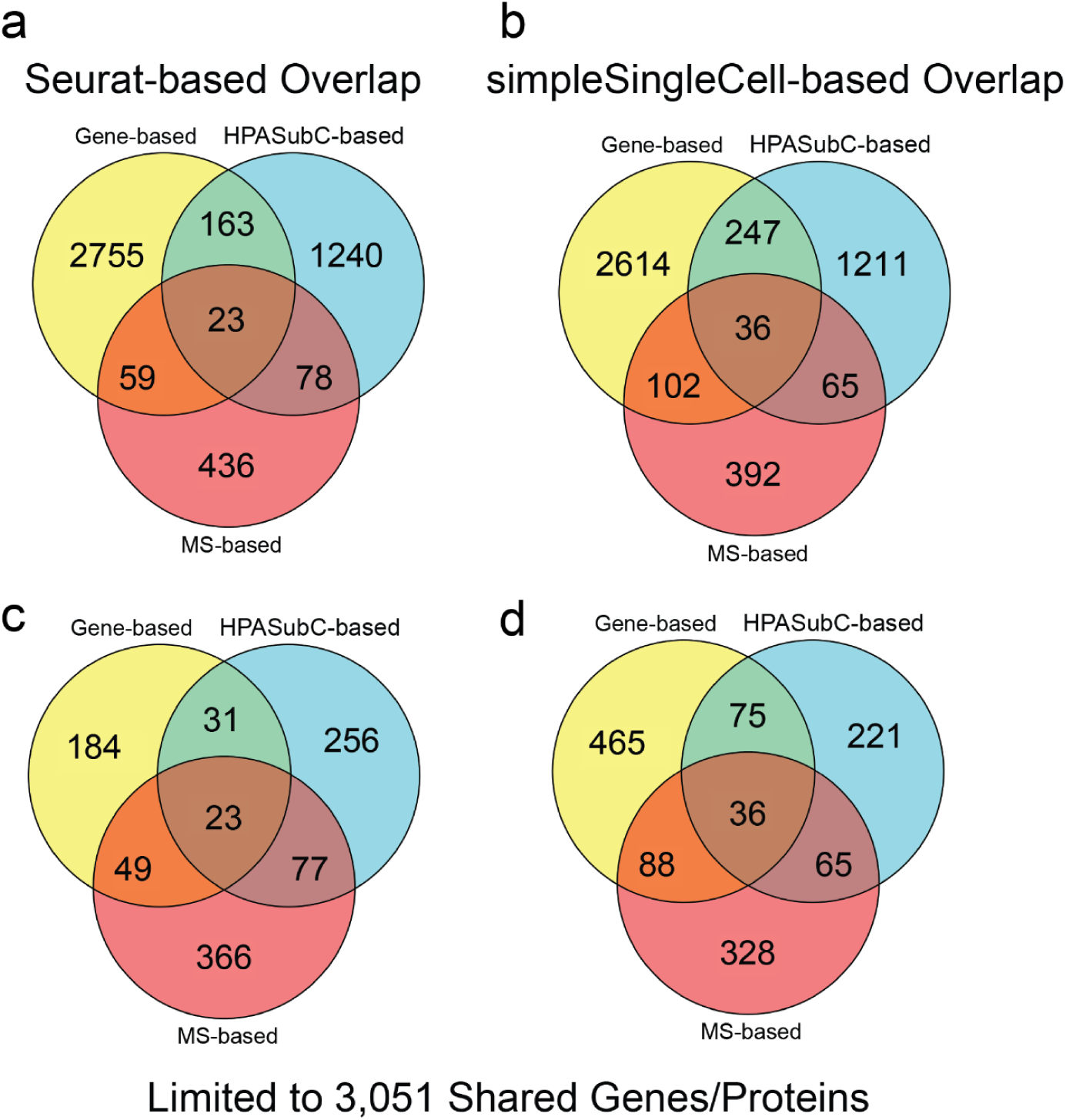
Venn diagrams comparing the three methods and two analysis types with the full datasets (top) and the limited datasets (below). **a)** A Seurat-based overlap including all mosaic genes/proteins. **b)** A simpleSingleCell-based overlap including all mosaic genes/proteins. **c** & **d)** Seurat and simpleSingleCell-based methods limited to the 3,051 genes/proteins shared across the three studies.

As so few genes were shared with the protein sets, we wondered if the computational approaches of the Seurat method limited the discovery of the correct variable genes. Therefore, we tried a second analysis approach, simpleSingleCell, to identify variable genes (21). By this method, there was an increase (N=36) in overlap of genes/proteins being identified by all three methods and more genes/proteins being identified by two methods (414) (Fig. 2b). Interestingly, comparisons limited to the shared gene/protein list resulted in the highest overlap between the MS- and gene-based datasets (Fig. 2d). A third method of using differential expression on the scRNA-seq data to compare the subset of 12 slow-twitch cells to all fast twitch (2X and 2A) or just fast 2X cells gave equivalent data to the simpleSingleCell approach.

### Shared, abundant transcripts by cell type

We then wondered about the extent to which highly abundant proteins/genes were driving our ability to detect mosaic proteins/genes. By normalized read counts of the scRNA-seq data, we determined the 50 most abundant transcripts by the average of each cell type in the three clusters determined by Seurat (Supplementary Data 2). Not surprisingly, the overall most abundant transcripts were *Ttn*, *Acta1* and *mt-Rnr2*. Of the 23 mosaic genes/proteins found by all three methods (using Seurat analysis), only *Myh1* and *Tnnt1* were on the list. Adding the mosaic genes from the simpleSingleCell analysis, seven additional genes (*Mylpf*, *Tnnt3*, *Tmp1*, *Tnni2*, *Eno3*, *Atp2a1* and *Pfkm*) were noted. This overall indicates that most abundant genes (≥41/50) are not consistently mosaic in skeletal myocytes.

### Species dichotomy in protein expression patterns

The generally low amount of overlap across the methods was unexpected. We wondered if this discrepancy particularly between the gene and protein data was the result of species differences in twitch type expression. To address this, we investigated staining patterns for three proteins. Two (DCAF11, ENO3) were selected as they had clear mosaic staining by human HPASubC images and no gene variation by Seurat analysis of the scRNA-seq. PVALB was selected for showing variation by the mouse scRNA-seq data, but no variation by HPASubC.

DCAF11 was robustly mosaic in human but non-mosaic in mouse. ENO3 was mosaic in both and PVALB was weakly mosaic in human but robustly mosaic in the mouse tissue (Fig. 3). This data suggested that discrepancies may relate to differences in mosaic protein expression between species (DCAF11) and possible technical causes (PVALB). Because ENO3 was mosaic in the mouse skeletal muscle, but not mosaic by Seurat gene expression analysis, we explored if a posttranscriptional form of regulation was occurring.

**Fig. 3.**
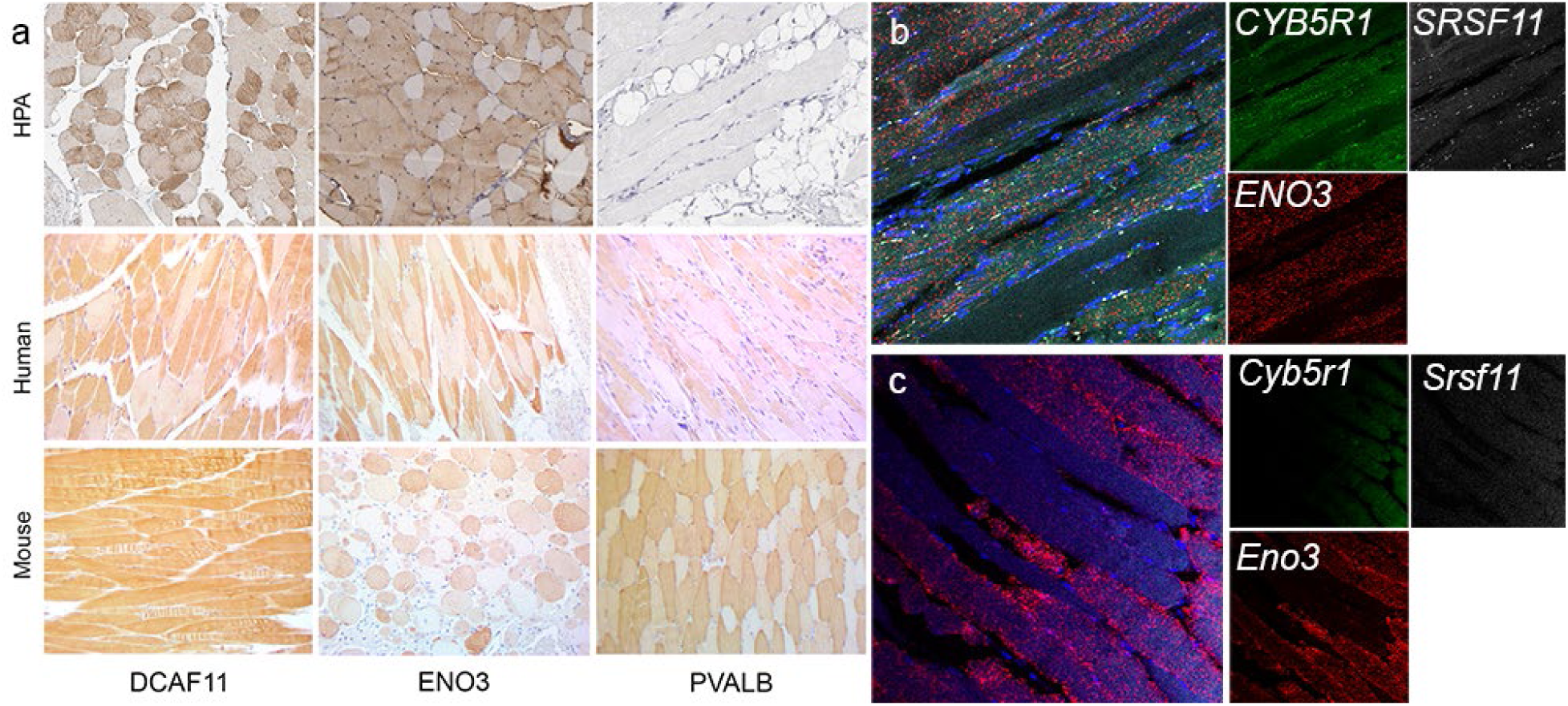
Representative IHC and RNA-ISH of discrepant proteins and genes. **a)** HPA images (top row) are mosaic for DCAF11 and ENO3 and negative for PVALB staining. Follow up staining validated the DCAF11 and ENO3 staining while suggesting a subtle mosaicism of PVALB in humans. In mice, ENO3 and PVALB are clearly mosaic, while DCAF11 is not. **b)** RNA-ISH demonstrates co-expression of *CYB5R1* and *ENO3* in a mosaic pattern. **c)** Only *Eno3* was observed (in a mosaic pattern) in mouse muscle by RNA-ISH.

### RNA-ISH indicates variable mosaicism

We performed RNA-ISH in both mouse and human skeletal muscles for *Eno3*, *Srsf11* and *Cyb5r1*. All of their protein products were mosaic by HPASubC and MS protein expression and had high or reasonably abundant gene expression (6552.8, 19.3, 201.5 pTPM respectively, HPA). None of these genes were variably mosaic in the mouse gene data. We found mosaic co-expression of all three genes in human skeletal muscle (Fig. 3). Whereas *ENO3* and *CYB5R1* RNA was diffusely present across human skeletal myocytes, *SRSF11* was localized to sub-cell membrane areas. In mouse muscle, *Eno3* was variably expressed, but neither *Cyb5r1* or *Srsf11* were identified, although their levels of expression (∼1,000x lower than *Eno3* in mouse) may be too low to be seen by this method.

### Many highly-supported variably expressed proteins were not previously identified

Thirty-six gene/proteins were variably expressed based on the simpleSingleCell, HPASubC and MS based analyses (Fig. 2d, Table 1). Of these, based on an extensive literature search, nine functionally diverse proteins are uniquely reported here as mosaic. Of the full 36, 22 were present in fast twitch myocytes and 14 in slow twitch myocytes based on the MS data. In addition to these 36, another 414 genes/proteins were identified by two complementary methods (Fig. 2). This includes well-known type specific proteins TNNC1 and TNNI1 (present in the HPASubC and simpleSingleCell datasets, but not variably expressed in the MS dataset).

**Table 1.** 37 Genes/Proteins identified as mosaic by all three methods based on simpleSingleCell analysis.

| Gene Name | Gene Symbol | Muscle Type- MS based | HPASubC Confidence | Seurat Norm. Counts | Mosaic Status |
| --- | --- | --- | --- | --- | --- |
| ArfGAP With Coiled-Coil, Ankyrin Repeat And PH Domains 2 | Acap2 | Fast | Likely | 1.79 | Unknown |
| Adenylate Kinase 1 | Ak1 | Fast | Likely | 5.75 | Known |
| Aldolase, Fructose-Bisphosphate A | Aldoa | Slow | Likely | 7.98 | Known |
| ATPase Sarcoplasmic/Endoplasmic Reticulum Ca2+ Transporting 1 | Atp2a1 | Fast | Real | 6.68 | Known |
| ATPase Sarcoplasmic/Endoplasmic Reticulum Ca2+ Transporting 2 | Atp2a2 | Slow | Real | 2.08 | Known |
| Calsequestrin 2 | Casq2 | Slow | Likely | 1.61 | Known |
| CD36 Molecule | Cd36 | Slow | Likely | 2.71 | Known |
| Creatine Kinase, Mitochondrial 2 | Ckmt2 | Slow | Likely | 4.54 | Known |
| DnaJ Heat Shock Protein Family (Hsp40) Member C3 | Dnajc3 | Fast | Likely | 2.2 | Unknown |
| Enolase 3 | Eno3 | Fast | Real | 6.97 | Known |
| ELKS/RAB6-interacting/CAST Family Member 1 | Erc1 | Fast | Likely | 3.14 | Unknown |
| Glyceraldehyde-3-Phosphate Dehydrogenase | Gapdh | Fast | Likely | 5.91 | Known |
| Glyoxalase Domain Containing 4 | Glod4 | Slow | Likely | 1.79 | Unknown |
| Glycerol-3-Phosphate Dehydrogenase 1 | Gpd1 | Fast | Likely | 3.22 | Known |
| Kinesin Family Member 5B | Kif5b | Fast | Likely | 3.83 | Unknown |
| Lactate Dehydrogenase A | Ldha | Fast | Real | 5.81 | Known |
| Lactate Dehydrogenase B | Ldhb | Slow | Real | 4.5 | Known |
| Myosin Binding Protein C, Fast Type | Mybpc2 | Fast | Real | 5.2 | Known |
| Myosin Heavy Chain 1 | Myh1 | Fast | Real | 7.91 | Known |
| Myosin Heavy Chain 6 | Myh6 | Slow | Likely | 1.79 | Known |
| Myosin Heavy Chain 8 | Myh8 | Fast | Likely | 5.41 | Known |
| Myosin Light Chain, Phosphorylatable, Fast Skeletal Muscle | Mylpf | Fast | Real | 3.47 | Known |
| Myosin Light Chain 3 | Myl3 | Slow | Real | 1.95 | Known |
| Myozenin 2 | Myoz2 | Slow | Real | 8.06 | Known |
| PDZ And LIM Domain 1 | Pdlim1 | Slow | Likely | 3.43 | Known |
| Peroxisomal Biogenesis Factor 19 | Pex19 | Fast | Likely | 2.41 | Unknown |
| Phosphofructokinase, Muscle | Pfkm | Fast | Likely | 2.2 | Known |
| Phosphoglycerate Kinase 1 | Pgk1 | Fast | Likely | 3.67 | Known |
| Ribosomal Protein S15a | Rps15a | Slow | Likely | 4.94 | Unknown |
| Thymosin Beta 4 X-Linked | Tmsb4x | Fast | Likely | 3.58 | Unknown |
| Troponin C2, Fast Skeletal Type | Tnnc2 | Fast | Real | 1.1 | Known |
| Troponin I2, Fast Skeletal Type | Tnni2 | Fast | Likely | 5.19 | Known |
| Troponin T1, Slow Skeletal Type | Tnnt1 | Slow | Real | 7.47 | Known |
| Troponin T3, Fast Skeletal Type | Tnnt3 | Fast | Real | 6.31 | Known |
| Topomyosin 1 | Tpm1 | Fast | Likely | 7.97 | Known |
| UDP-Glucose 6-Dehydrogenase | Ugdh | Slow | Likely | 8.01 | Unknown |

Finally, another 4,217 genes/proteins were variably expressed by one method. Of this group, 1,211 were detected by HPASubC and 270 of these proteins were scored as “real” with clear patterns of mosaicism (Supplementary Fig. 4).

## Discussion

We describe the first proteogenomic analysis of skeletal muscle single fiber types using combined scRNA-seq, spatial proteomics, and MS proteomics. Because delineations of skeletal muscle fiber types are known and this project was exclusive to this one cell type, our study is a useful model system to evaluate combining and synthesizing gene and protein data into a coherent description of a cell. Also, by utilizing a deep sequencing approach and fewer cells, we were not limited to just classifying a cell, but rather had sufficient data to delve into full gene expression. Our data identifies common themes across the methods, but also significant differences and complexities in gene/protein assignments.

**Supplementary Figure 4.**
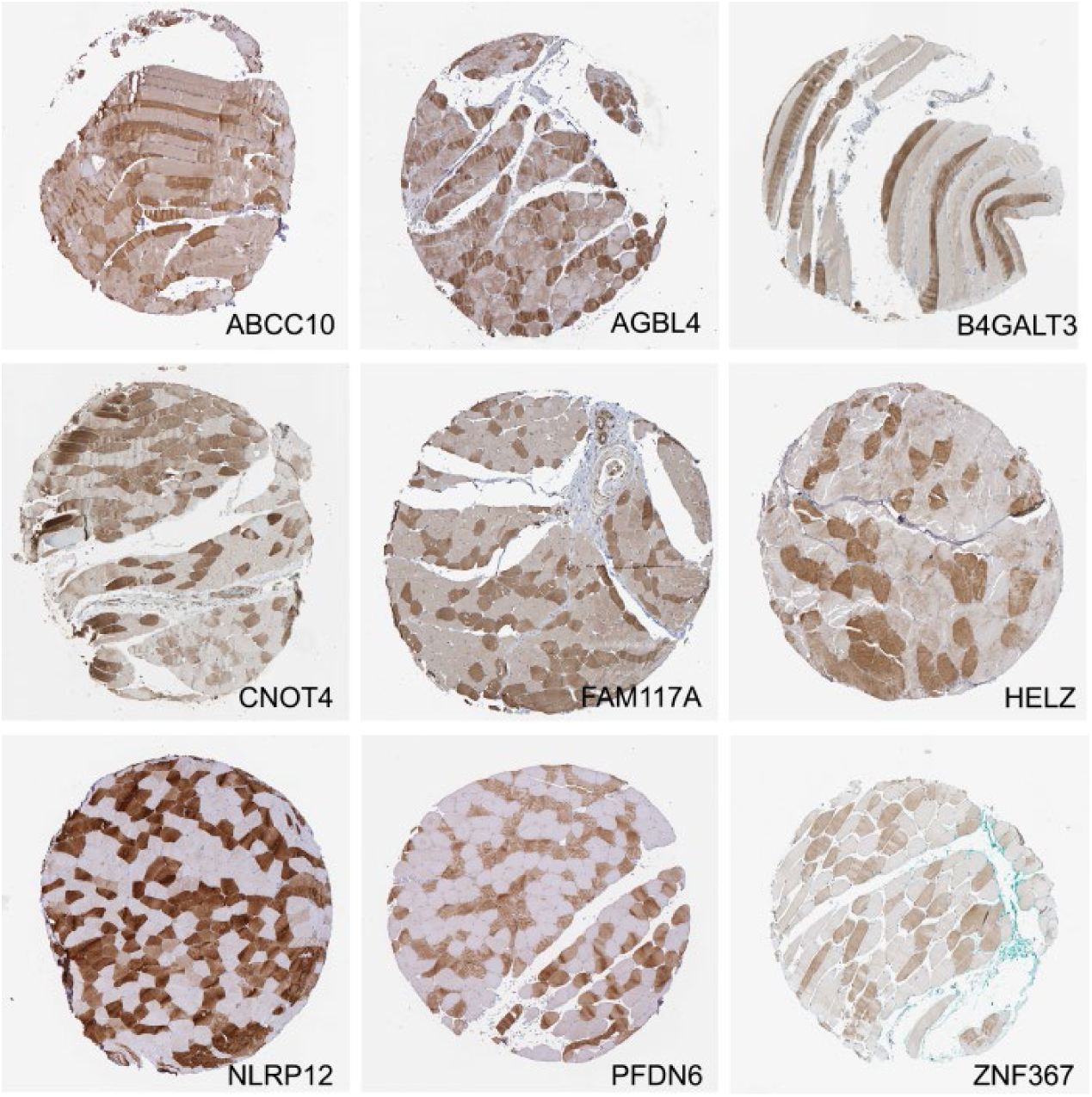
Nine representative images of 270 proteins scored as real mosaicism using HPASubC, but not identified by other methods. All images from HPA.

Regardless of the method and genes/proteins used to cluster, we found general agreement on the major types of skeletal muscle myocytes. We identified a small group of slow twitch cells that clustered with fast 2A cells. These groups were consistently clustered away from two clusters of fast 2X cells. The differences between these two fast 2X groups, described herein as 2X_a_ and 2X_b_, are open to interpretation. The simplest explanation is that some axonal material remained variably adherent to skeletal muscle cells through the NMJ, and these 20+ genes resulted in the separation observed by UMAP (Fig. 1a). This would imply a technical cause of the two fast 2X cell subtypes as a result of myocyte isolation. Adherent cell fragments are likely to be a global issue for some cell type isolation, although it would not impact nuclear scRNA-seq studies. A more interesting explanation is variable neuronal transfer of mRNAs across the NMJ into the skeletal muscles via extracellular vesicles (22, 23). This would imply a real state-difference in these cells, notable only by the deep sequencing strategy employed. Regardless, of which is accurate, this division is unlikely to indicate true separate fast 2X subtypes. In fact, the cross-referenced proteomic data was useful in demonstrating the arbitrary nature of this delineation (Fig. 1i).

The extent of overlap of mosaic genes/proteins across the methods was surprisingly low. Only 36 genes/proteins were cross-validated across all three approaches using the simpleSingleCell method (Table 1). This list included well-known, fiber-type specific proteins such as MYH1 and MYH6 and newly described mosaic proteins like DNAJC3 and GLOD4. The lack of agreement across methods has made it difficult to confidently state how many proteins/genes are variable by twitch pattern and further demonstrates the challenge of relying on a single method. If a gene or protein is mosaic by two methods, this number climbs to 450. If all mosaic genes and proteins are included, this increased to >4,500 genes/proteins. Over 1,600 proteins appear to be mosaic by the HPASubC method alone (Supplementary Data 1 and Supplementary Fig. 4).

The reason for the variability in mosaic genes/proteins is certainly multifactorial. One potential major difference is the comparison across two separate species (mouse and human). As we noted with the DCAF11 IHC, this protein was mosaic in human muscle but did not appear to be mosaic in mouse. Secondly, some genes have markedly variable expression levels between the two species. While *CYB5R1* and *SRSF11* are robustly expressed in human muscle at 201.5 and 19.3 pTPM (in HPA), they were only 17.7 and 1.9 FPKM in our mouse scRNA-seq. It is also possible that post-transcriptional regulation leads to more extreme expression variation in proteins than genes. As described above, extreme expression dichotomy in *Myh* genes was less than in similar MYH protein data (Fig. 1d) (5).

Our study represents the first use of LP-FACS to isolate single myofibers for scRNA-seq. As skeletal myocytes are often long, stretching across the length of a muscle, isolation techniques (particularly from human samples) may rely on the use of biopsies or otherwise fragmented myocytes. To test the effect of myocyte fragmentation on scRNA-seq data quality, we used a liberal gating strategy of our dissociated myocytes (including both EXT-high/TOF-low and EXT-high/TOF-high populations) as well as directly sequencing fragmented myocytes generated through a pseudo-biopsy approach. Disappointingly, we found that a large portion of our sequenced myocytes were of poor quality, including those from our pseudo-biopsy approach. By contrast, the highest quality data likely came from fully intact myocytes, in particular the EXT-high/TOF-high population. Because this population is almost completely enriched for intact myocytes, we believe that future experiments using LP-FACS to isolate skeletal myocytes should focus solely on the EXT-high/TOF-high population. We are confident that this will allow for a much higher percentage of good quality scRNA-seq libraries, akin to what we have observed previously with LP-FACS isolation of cardiac myocytes (17). These results also mean that more work must be done to identify better isolation methods for human skeletal muscle. Current methods of human skeletal muscle biopsying from the quadriceps only obtains muscle fragments and thus more creative methods to obtain full length fibers or non-damaged fibers must be considered.

Technical factors also impact our ability to detect mosaicism on all platforms. Discovery mass spectrometry is challenged to identify low abundance proteins. Having low input from single fibers was further limiting and reduced the ability to computationally distinguish expression differences in low abundance proteins. Most fibers had between 500-700 proteins identified. As we have stated repeatedly, IHC in the HPA is subject to false positive staining from shared epitopes (20, 24–26). It also incurs false negative staining for failed antibodies or antibodies with staining parameters designed for other tissues. Further, some genes/proteins observed in the other datasets were missing from the HPA data. The gene data was also limited in the number of total cells analyzed (171) and the rarity of slow twitch cells from this muscle. Cross-cell contamination, may have also stunted the differences between cell types (27).

In conclusion, we have created the first proteogenomic analysis of gene/protein mosaicism in skeletal muscle. We replicated the known fiber types of slow, fast 2A, and fast 2X, as well as greatly expanded our understanding of genes and with variable expression across these cell types.

## Methods

### Isolation and Sequencing of Adult Skeletal Myocytes

Experiments were performed using C57BL/6J mice greater than 3 months of age. To isolate skeletal myocytes, we performed collagenase-based digestion of the flexor digitorum brevis (FDB), a short muscle of the hind feet, as per previously established protocols (28). We tested two separate approaches to isolating myocytes. In the first approach, we dissected the FDB from tendon to tendon prior to digestion, enabling isolation of fully intact myocytes. In the second approach, we cut small portions of the FDB muscle using scissors. We reasoned that the latter approach would broadly mimic skeletal muscle biopsy as might be done, for example, from a human patient sample. The FDB was transferred to a dish containing DMEM with 1% penicillin/streptomycin, 1% fetal bovine serum, and 2mg/mL Collagenase Type II (Worthington). Muscle was digested for 1.5 hours in a 37C cell incubator with 5% CO_2_. Subsequently, the muscle was transferred to a dish containing media without collagenase, and gently triturated to release single myocytes. Large undigested chunks and tendons were removed with tweezers prior to single cell isolation.

We subsequently isolated single myocytes through large particle fluorescent-activated cell sorting (LP-FACS), using a flow channel size of 500 µm. The COPAS SELECT Flow Pilot Platform (Union Biometrica) was employed. Using time-of-flight (TOF, measuring axial length) and optical extinction (EXT, measuring optical density) parameters, we found that skeletal myocytes separated into three populations – an EXT-low population, EXT-high/TOF-low population, and EXT-high/TOF-high population (Supplementary Fig. 1A). The EXT-high/TOF-high population was comprised almost entirely of intact myofibers with lengths > 400 µm, suggesting successful sorting of large myocytes (Supplementary Fig. 1B). Interestingly, the EXT-high/TOF-low population was composed of what appeared to be rod-shaped fragments that maintained sarcomeric proteins, albeit disrupted (Supplementary Fig. 1C). The EXT-low population was comprised mostly of debris and dead cells, as previously observed with cardiac myocytes (Supplementary Fig. 1D). The EXT-high/TOF-low population qualitatively resembled our pseudo-biopsy isolated myocyte fragments (Supplementary Fig. 1E), which also shared similar TOF and EXT parameters (not shown). To our knowledge, this is the first FACS-based single cell RNA-seq study of skeletal myocytes; thus, we adopted a broad gating strategy for isolation of single cells. We sorted 700 EXT-high myocytes (comprised of both TOF-high and TOF-low populations) as well as 100 myocyte fragments isolated through the pseudo-biopsy method.

These sorted cells were placed individually into 96-well plates. Capture plate wells contained 5 µl of capture solution (1:500 Phusion High-Fidelity Reaction Buffer, New England Biolabs; 1:250 RnaseOUT Ribonuclease Inhibitor, Invitrogen). Single cell libraries were then prepared using the previously described mcSCRB-seq protocol (18, 19). Briefly, cells were subjected to proteinase K treatment followed by RNA desiccation to reduce the reaction volume. RNA was subsequently reverse transcribed using a custom template-switching primer as well as a barcoded adapter primer. The customized mcSCRB-seq barcode primers contain a unique 6 base pair cell-specific barcode as well as a 10 base pair unique molecular identifier (UMI). Transcribed products were pooled and concentrated, with unincorporated barcode primers subsequently digested using Exonuclease I treatment. cDNA was PCR-amplified using Terra PCR Direct Polymerase (Takara Bio). Final libraries were prepared using 1ng of cDNA per library with the Nextera XT kit (Illumina) using a custom P5 primer as previously described.

### scRNA-seq sequencing and analysis

Pooled libraries were sequenced on two high-output lanes of the Illumina NextSeq500 with a 16 base pair barcode read, 8 base pair i7 index read, and a 66 base pair cDNA read design. To analyze sequencing data, reads were mapped and counted using zUMIs 2.2.3 with default settings and barcodes provided as a list (29). zUMIs utilizes STAR (2.5.4b) (30) to map reads to an input reference genome and featureCounts through Rsubread (1.28.1) to tabulate counts and UMI tables (30, 31). Reads were mapped to the mm10 version of the mouse genome. We used GRCm38 from Ensembl concatenated with ERCC spike-in references for the reference genome and gene annotations. Dimensionality reduction and cluster analysis were performed with Seurat (2.3.4) (32).

### Seurat and simpleSingleCell

Analysis was performed using the Seurat R toolkit V3.1.1 for this dataset (32). Initial filtering removed lower quality cells (read count <5000 RNAs detected or >20% mitochondrial genes) before sctransform normalization (33). A standard Seurat workflow was initially used for data analysis. This workflow identifies a subset of genes with high cell-to-cell variation within the scRNA-seq data. This subset is subsequently used as input to principal component analysis as well as downstream nonlinear dimensionality reduction methods such as Uniform Manifold Approximation and Projection (UMAP). Additionally, Seurat also allows for use of custom gene lists as input to downstream analysis. This allowed us to use two custom gene lists, specifically those derived from orthologous genes to mosaic proteins in the visual (HPASubC) dataset (20) or the differentially expressed proteins in the MS proteomic dataset (5). Thus each of our three gene lists, one produced by Seurat’s workflow, another visual proteomic-based gene list, and a final mass spectrometry-based gene list defining known muscle cell types, were used one at a time to subset our initial data set and generate principal components for downstream analysis.

After determining clustering via these three approaches, UMAPs were generated alongside with heat maps representing the top genes in clusters as determined by each gene set used for PCA. Overlapping genes between the HPAsubC data, MS data, and significant genes determined by Seurat were also examined for overlaps. Gene expression for *Trdh, Lama4, Ryr1, Dpp10, Pde4dip, Sugct,* and *Myh1* was plotted across two fast 2X_a_ clusters based on the HPASubC data.

### Simple Single Cell and Scran

Simple single cell 1.8.0 workflow was followed using scran 1.12.1 for normalization of raw counts and fitting a mean-dependent trend to the gene-specific variances in single-cell RNA-seq data (21). In line with this, we decomposed the gene-specific variance into biological and technical components and selected the top 3000 genes for comparisons.

### RNA-ISH

Mouse and human skeletal muscles were obtained at necropsy (>3 month old) and rapid autopsy (66 year old male), the latter under an IRB-approved protocol. Tissues were immediately fixed in formalin and paraffin-embedded blocks were created, from which 5 micron slides were made.

Custom probes for RNA *in situ* hybridization (RISH) were obtained from RNAscope (ACDBio). These probes were designed to detect human and mouse forms of the following genes: *ENO3* (GenBank accession nm_001976.5), *CYB5R1* (nm_016243.3), *SRSF11* (nm_004768.5), *Eno3* (nm_007933.3), *Cyb5r1* (nm_028057.3), and *Srsf11* (nm_001093753.2). Each probe set targeted all validated NCBI refseq transcript variants of the gene.

The Multiplex Fluorescent Reagent Kit v2 (ACDBio) was used following the manufacturer’s instructions. Briefly, FFPE tissue slides were baked for one hour at 60°C. The slides were subsequently deparaffinized with xylene, rinsed with 100% ethanol and air-dried. After application of hydrogen peroxide and washing, slides were treated with target retrieval reagent in a steamer (>99°C) for 20 minutes. Then, the tissue was permeabilized using a protease.

Hybridization of the probes to the targeted mRNAs was performed by incubation in a 40°C oven for 2 hours. After washes, the slides were processed for the standard signal amplification and application of fluorescent dye (Opal dye 520, 570 and 620, AKOYA Biosciences) steps. Finally, the slides were counterstained with DIPA, mounted with Prolong Gold Antifade Mounting solution (Invitrogen) and stored in a 4°C room. The fluorescent images were obtained in the Johns Hopkins Microscope Core Facility using a Zeiss LSM700 Laser scanning confocal microscope.

### Immunohistochemistry

The same tissues described above were used for standard immunohistochemistry. Antibodies were obtained for WDR23/DCAF11 (bs-8388R, Bioss Antibodies), PVALB (A2781, Abclonal), and ENO3 (ARP48203_T100, Aviva Systems Biology) that were reported to cross react to both human and mouse. Immunohistochemistry was performed as described previously (25, 34).

### HPA and HPASubC

The HPA is a comprehensive repository of IHC stained tissue microarrays for numerous tissues, including skeletal muscle (35, 36). The HPASubC tool can rapidly and agnostically interrogate images of the HPA to characterize specific staining patterns in organs (20, 24, 26). HPASubC v1.2.4 was used to download 50,351 skeletal muscle tissue microarray images covering 10,301 unique proteins from the HPA website (v18). The images were individually reviewed using HPASubC by K.M.F to evaluate the presence of a mosaic pattern of protein expression based on IHC staining. The classification of mosaicism was based on a pre-study training set of 300 images from HPA reviewed collaboratively (K.M.F and M.K.H). Mosaicism was defined as a dispersed pattern of differential staining in which a significant number of non-adjacent muscle fibers had a higher staining intensity than the surrounding fibers, preferably persisting across the entire microarray. All positive selections made by the trainee were reviewed and rescored, as needed, by a board-certified pathologist (M.K.H.).

After an initial fast review of the images, a re-review to score the images was performed. A three-tiered classification system was used indicating increasing certainty of mosaicism: 0 indicated the absence of mosaic staining; 1 indicated unknown mosaic staining; 2 indicated likely mosaic staining; 3 indicated real mosaic staining. Scoring evaluation was based on the quality of the mosaic pattern, including stain intensity differential between fibers, the presence of “blush”/incomplete staining within cells, and the consistency and completeness of the fiber staining pattern throughout the sample. HPASubC was used on an Apple MacBook Pro running macOS Sierra v10.12.6 with 8 GB RAM and 3.1 GHz CPU and a Dell Precision Tower 3620 running Windows 10 with 16 GB RMA and a 3.7 GHz CPU.

### Conversion of gene and protein symbols

To identify orthologs across human and mouse genes/proteins we had to synchronize across gene/protein names and across the species. We used the David Gene ID Conversion Tool (https://david.ncifcrf.gov/conversion.jsp), BioMart at Ensembl (http://useast.ensembl.org/biomart/martview/e8a4fba4cb5c0be7a30841471b55674d), UniProt Retreive/ID mapping (https://www.uniprot.org/uploadlists/) and direct searches at both UniProt and GeneCards (https://www.genecards.org/), to cross integrate the human protein symbols, mouse gene symbols, human gene symbols and ENSG IDs (37–39).

### Gene Ontology (GO) Validation

GO was performed on the 557 most variable genes between two fast 2X clusters (2X_a_ and 2X_b_) using the Gene Ontology resource (http://geneontology.org/) and selecting for cellular component.

### Mass Spectrometry (MS) Data Set

We utilized the Murgia et al. human skeletal muscle fiber MS-based proteomic dataset (5). This contained information from 3,585 proteins across 152 fibers from 8 donors (5). The ratio of expression of proteins between Type 1 and Type 2A cells were determined using Table S6 of Murgia et al. Five hundred and ninety-six proteins with >2.3 fold differences between cell types were selected. Label-free quantification (LFQ) data, from Supplemental Table S4, for the 154 human single muscle fiber proteomics was obtained. The log2 transformed LFQ data was converted to raw values and only proteins expressed across all fiber types (n=94) were considered for plotting UMAP as described (5). Functions of the R-package Seurat (Version 3.1.1) were executed sequentially to derive a UMAP along with its dependency library “uwot (Version 0.1.4)” in R (Version 3.6.1) (40, 41). A Seurat object of the data matrix was created using ‘CreateSeuratObject’ with default parameters. This data was normalized using the ‘NormalizeData’ function and outlier proteins were identified using the ‘FindVariableFeatures.’ Proteins across the fiber types were scaled and centered to create a PCA object using ‘ScaleData’ and ‘RunPCA’ respectively. Further, k-nearest neighbors and shared nearest neighbor for each fiber type were generated on the Seurat object using ‘FindNeighbors’ and ‘FindClusters’to plot UMAP using ‘RunUMAP’. All of these functions were executed using default parameters. The clustering obtained with UMAP was overlaid with the classification of muscle fiber types based on Murgia et al. using ggplot2 (Version 3.2.1).

## Data availability

Mouse skeletal muscle sequencing was deposited at the Sequence Read Archive (SRA – SRP241908) and the Gene Expression Omnibus (GSE143636).

## Code availability

All analysis scripts are available at GitHub (https://github.com/mhalushka/Skeletal_muscle_mosaicism).

## Acknowledgements

The authors thank Efrain Ribeiro for his helpful comments on the project. M.K.H. was supported by grants 1R01HL137811, R01GM130564, and P30CA006973 from the National Institutes of Health and 17GRNT33670405 from the American Heart Association. T.O.N. was supported by grant R01GM130564. M.N.M. was supported by R01HL137811 and the University of Rochester CTSA award number UL1TR002001. A.Z.R was supported by R01GM130564. C.K. and S.K. were supported by NIH R01HD086026, TEDCO 2019-MSCRFD-5044, and the JHU Discovery Award. S.K. was supported by fellowship 20PRE35200028 from the American Heart Association.

## Contributions

K.M.F helped conceive the project and generated proteomic data. S.K. and B.L.L. generated the skeletal muscle sequencing library. X.Y and K.F-T. performed IHC and RISH. R.X.V., S.K., T.O.N, A.H.P. and M.N.M. performed analysis. C.K. and D.A.K. oversaw the library preparation. A.Z.R helped develop the project. M.K.H. conceived the project, performed analyses and wrote the manuscript. All authors contributed toward revisions of the manuscript.

## Conflicts of interest

The authors declare no conflicts of interest.

## Notes

#### Summary of Updates

The first name of the first author was missing the terminal 'e'. No other changes were made.

https://github.com/mhalushka/Skeletal_muscle_mosaicism

